# Multiple DNA repair pathways prevent acetaldehyde-induced mutagenesis in yeast

**DOI:** 10.1101/2024.01.07.574575

**Authors:** Latarsha Porcher, Sriram Vijayraghavan, James McCollum, Piotr A Mieczkowski, Natalie Saini

## Abstract

Acetaldehyde is the primary metabolite of alcohol and is present in many environmental sources including tobacco smoke. Acetaldehyde is genotoxic, whereby it can form DNA adducts and lead to mutagenesis. Individuals with defects in acetaldehyde clearance pathways have increased susceptibility to alcohol-associated cancers. Moreover, a mutation signature specific to acetaldehyde exposure is widespread in alcohol and smoking-associated cancers. However, the pathways that repair acetaldehyde-induced DNA damage and thus prevent mutagenesis are vaguely understood. Here, we used *Saccharomyces cerevisiae* to systematically delete genes in each of the major DNA repair pathways to identify those that alter acetaldehyde-induced mutagenesis. We found that deletion of the nucleotide excision repair (NER) genes, *RAD1* or *RAD14*, led to an increase in mutagenesis upon acetaldehyde exposure. Acetaldehyde-induced mutations were dependent on translesion synthesis as well as DNA inter-strand crosslink (ICL) repair in *Δrad1* strains. Moreover, whole genome sequencing of the mutated isolates demonstrated an increase in C→A changes coupled with an enrichment of gCn→A changes in the acetaldehyde-treated *Δrad1* isolates. The gCn→A mutation signature has been shown to be diagnostic of acetaldehyde exposure in yeast and in human cancers. We also demonstrated that the deletion of the two DNA-protein crosslink (DPC) repair proteases, *WSS1* and *DDI1*, also led to increased acetaldehyde-induced mutagenesis. Defects in base excision repair (BER) led to a mild increase in mutagenesis, while defects in mismatch repair (MMR), homologous recombination repair (HR) and post replicative repair pathways did not impact mutagenesis upon acetaldehyde exposure. Our results in yeast were further corroborated upon analysis of whole exome sequenced liver cancers, wherein, tumors with defects in ERCC1 and ERCC4 (NER), FANCD2 (ICL repair) or SPRTN (DPC repair) carried a higher gCn→A mutation load than tumors with no deleterious mutations in these genes. Our findings demonstrate that multiple DNA repair pathways protect against acetaldehyde-induced mutagenesis.

## Introduction

Acetaldehyde is produced as a metabolic byproduct in response to numerous exogenous agents, including tobacco smoke, air pollution, food products, and is a major product of alcohol metabolism [1, 2]. Individuals who are unable to metabolize acetaldehyde because of mutations in the aldehyde dehydrogenase 2 gene (*ALDH2*) have an elevated risk for alcohol- or smoking-related carcinogenesis [3-6]. Mice with defects in *ALHD2* and the Fanconi Anemia pathway also demonstrate increased predisposition to leukemia [7]. The US Environmental Protection Agency currently classifies acetaldehyde as a group B2 carcinogen [8, 9].

Genome instability is the primary mechanism by which acetaldehyde exposure has been hypothesized to promote carcinogenesis. Biochemically, acetaldehyde can directly form a variety of DNA lesions including N^2^-ethylidene-2′-deoxyguanosine (N^2^-Eti-dG), 1,N^2^-propano-2′-deoxyguanosine, and G-G inter- and intra-strand crosslinks [10-14]. Alcohol consumption and tobacco smoking lead to a strong increase in the levels of N2-Eti-dG in the oral cavity and in blood [15, 16].

Interestingly, despite the increased DNA adducts and DNA damage seen with acetaldehyde exposure, studies in yeast [17] and in human induced pluripotent stem cells [18] determined that acetaldehyde was not mutagenic. Contrary to these studies, we recently demonstrated that acetaldehyde is a single stranded DNA mutagen in yeast [19]. Acetaldehyde exposure was demonstrated to produce C→A mutations in an gCn context (mutated residue is capitalized and n denotes all bases) in yeast strains. Such mutagenesis was found to be dependent on the presence of single stranded DNA in yeast wherein DNA repair was not functional [19]. This gCn→A signature was further identified in tumors associated with smoking or alcohol consumption in whole-genome-sequenced cohorts from the Pan-cancer Analysis of Whole Genomes (PCAWG) [20] and whole-exome-sequenced cohorts from the International Cancer Genome Consortium (ICGC) [19, 21]. While our findings indicate widespread acetaldehyde-induced mutagenesis, it is likely that substrate limitations and highly efficient DNA repair constrain the detection of mutations.

Several studies have highlighted the DNA repair pathways that render cells more susceptible to the effects of acetaldehyde exposure. Mice deficient in ALDH2 and FAND2 had a higher mutation load in hematopoietic stem cells due to accumulation of endogenous DNA damage. However, their mutation pattern was the same as wild-type mice [22]. In addition, activity of FANCD2 in resolving inter-strand crosslinks (ICLs) was implicated in avoidance of acetaldehyde-induced genotoxicity in oral keratinocytes [23]. *ALDH1B1* and *MSH2* deletion in mice also led to increased colonic tumors [24]. The Fanconi anemia pathway was also found to be responsible for repair of DNA intra-strand crosslinks created *in vitro* on plasmids [25]. Cell extracts deficient in xeroderma pigmentosum group A (XPA) proteins or human cell lines with XPA defects were unable to repair acetaldehyde-induced DNA damage on plasmid DNA, indicating a role of NER in repair of acetaldehyde DNA damage [26]. Additionally, in *Schizosaccharomyces pombe,* defects in homologous recombination, aldehyde clearance pathways, nucleotide excision repair, fork protection complex and checkpoint response pathways were found to sensitize cells to exogenous acetaldehyde [27]. However, because these studies primarily focused on DNA damage and cell death as the primary outcome of acetaldehyde exposure as opposed to mutagenesis, by and large, the genetic determinants of acetaldehyde-induced mutagenesis are poorly understood.

Here, we systematically tested deletions in genes within all the major DNA repair pathways to identify those pathways that function to prevent mutagenesis upon acetaldehyde exposure. Similar to previous works, we determined that acetaldehyde exposure is not mutagenic in DNA repair proficient yeast. However, ablation of nucleotide excision repair (NER) and DNA-protein crosslink (DPC) repair pathways lead to an increase in acetaldehyde mutagenesis in yeast. We further demonstrate that increased mutagenesis in NER-deficient strains was dependent on the activity of polymerases involved in translesion synthesis (TLS). Interestingly, deletion of *PSO2* in an NER-deficient genetic background lowered acetaldehyde-induced mutagenesis. These data point at a role of DNA inter-strand crosslink (ICL) repair pathway in acetaldehyde-mediated mutagenesis. Additionally, we uncovered a role of base excision repair (BER) in acetaldehyde-induced mutagenesis, with the loss of *APN1* and *APN2* leading to a mild increase in acetaldehyde mutagenesis. Interestingly, while homologous recombination and mismatch repair have previously been implicated in the repair of acetaldehyde-induced DNA damage, these two pathways do not play a role in acetaldehyde-induced mutagenesis in our study. Overall, our work demonstrates that multiple DNA repair pathways can efficiently remove DNA damage induced by acetaldehyde, thus preventing mutagenesis.

## Results

### Acetaldehyde is not mutagenic in DNA repair proficient yeast strains

We previously demonstrated that acetaldehyde was highly mutagenic to single-stranded DNA in yeast [19]. However, various studies in bacteria, yeast and iPSCs showed that acetaldehyde exposure was not mutagenic to cells [17, 18, 28]. To address this discrepancy in observations, we grew haploid yeast cultures overnight and sub-cultured into fresh media for 3 hours to obtain actively dividing yeast strains (Fig 1A). We hypothesized that yeast cultures with active DNA replication and transcription should have single stranded DNA available for acetaldehyde-induced mutagenesis. We incubated yeast cultures with varying concentrations of acetaldehyde and and asked whether there was an impact on viability and/or mutagenesis using plating assays (Fig 1A). We did not note observe any increase in Can^R^ mutation frequencies in the wild-type isolates treated with the different concentrations of acetaldehyde. Also, no changes in cellular viability were observed at 1% acetaldehyde as compared to the water treated samples. These data corroborate the findings from others and demonstrate that acetaldehyde is likely not mutagenic to yeast with proficient DNA repair pathways (Fig 1B,C,D, Table S1 and S2).

**Figure 1.**
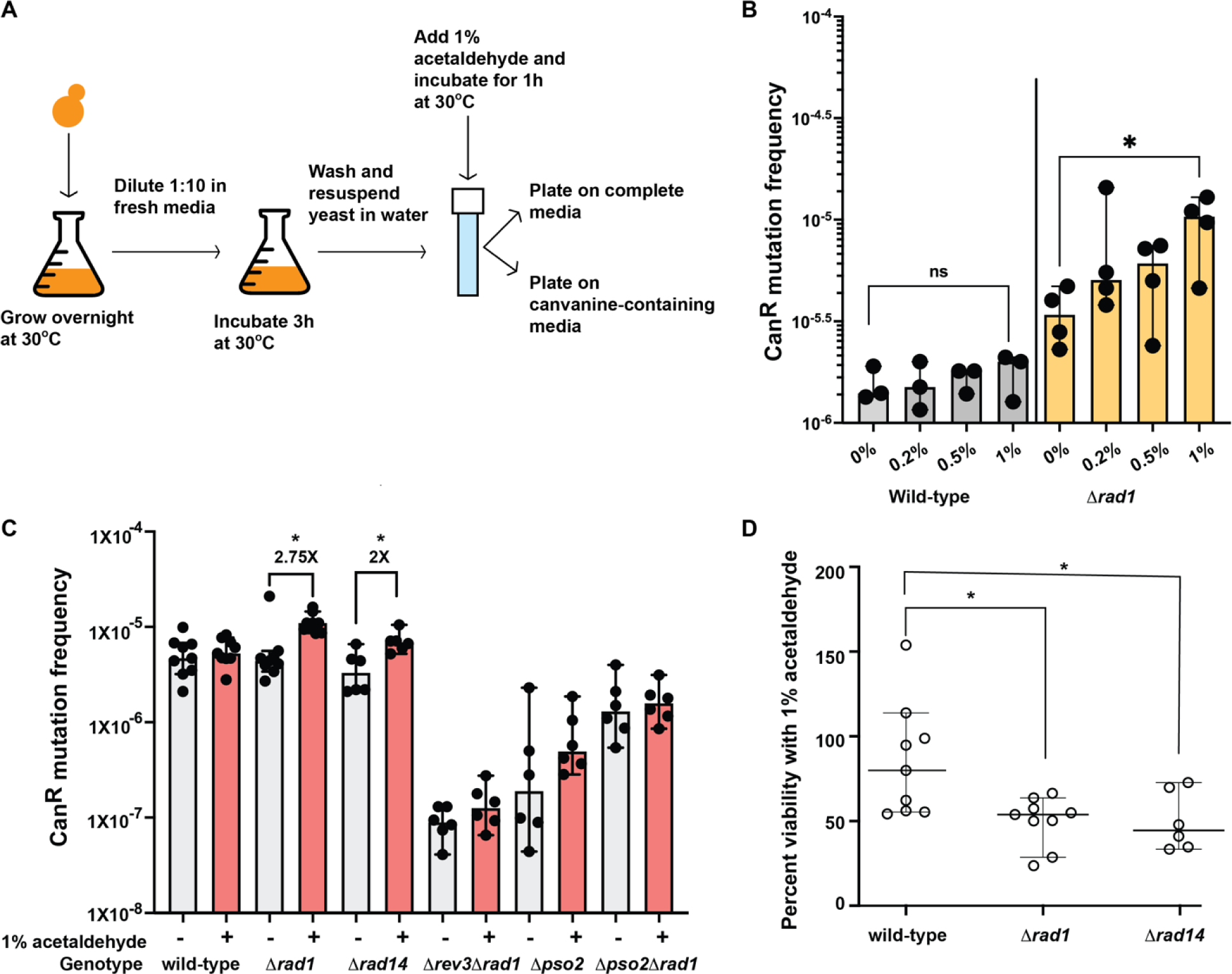
Nucleotide excision repair prevents acetaldehyde-induced mutagenesis in yeast. A) Schematics of the experiment. Yeast cultures were grown overnight in liquid media. Cultures were diluted and allowed to grow in fresh media for 3 hours. Cultures were then spun down, washed, and resuspended in water and moved to a screw-cap tube. Acetaldehyde was added and the cultures were incubated for 1 hour. Appropriate dilutions were plated on complete and canavanine containing media to score for mutations in the *CAN1* gene. B) Mutation frequencies of wild-type and *Δrad1* strains treated with different concentrations of acetaldehyde. Bars indicate median mutation frequency and error-bars denote range. C) Mutation frequencies of yeast strains treated with no mutagen (-) or 1% acetaldehyde (+). Bars indicate median mutation frequencies and error bars indicate 95% confidence intervals. D) Quantitative survival assay of yeast strains treated with acetaldehyde as compared to no mutagen. Median and 95% confidence intervals are depicted. P-values were calculated using a one-tailed Mann-Whitney U-test with the hypothesis that acetaldehyde-treated samples have increased mutation frequencies. * denotes P-values < 0.05.

### Defects in nucleotide excision repair increases acetaldehyde-induced mutation frequencies

Rad1 (ERCC4 in humans) is the endonuclease involved in NER (reviewed in [29]). We treated *Δrad1* isolates with varying concentrations of acetaldehyde as described above (Fig 1) and saw a significant increase in mutagenesis in the *Δrad1* cultures treated with 1% acetaldehyde for 1 hour as compared to the untreated cultures (2.75-fold increase, P-value using one sided Mann-Whitney U test = 0.002) (Fig 1B, C). Additionally, we noted a decrease in cellular viability in these isolates upon acetaldehyde exposure (median viability 43%), indicating that NER is a major pathway for repair of acetaldehyde-induced DNA damage (Fig 1C, D, Tables S1 and S2).

To further validate our findings on the importance of NER in repairing acetaldehyde-induced DNA damage, we deleted the *RAD14* gene and analyzed mutation frequencies upon acetaldehyde exposure. Rad14 (XPA in humans) recognizes and binds DNA damage to promote NER [29]. *Δrad14* strains treated with acetaldehyde demonstrated a similar increase in mutagenesis as compared to the untreated cultures (2-fold increase, P-value using one sided Mann-Whitney U Test = 0.0043) (Fig 1C, Table S1). Acetaldehyde treatment led to a reduction in viability of *Δrad14* cultures to 45% as compared to the untreated cultures (Fig 1C, D, Table S2).

Previously, we also demonstrated that acetaldehyde-adducts in single-stranded DNA are bypassed by TLS polymerases leading to mutagenesis [19]. Rev3 is the catalytic subunit of DNA Polζ and is essential for TLS in yeast. To determine if the increase in mutations seen in the *Δrad1* strains upon treatment with acetaldehyde was dependent on TLS, we deleted *REV3* in the *Δrad1* strain. We found that endogenous level of mutagenesis was drastically reduced in the *Δrad1Δrev3* isolates compared to the wild-type isolates. Also, we did not see any increased mutagenesis upon treatment of these cultures with acetaldehyde indicating that Rev3 was required for acetaldehyde-induced mutagenesis in the *Δrad1* strains (Fig 1C, Table S1). Our data demonstrates that unrepaired acetaldehyde-induced DNA adducts in the NER-deficient *Δrad1* isolates are bypassed by TLS resulting in mutagenesis.

### DNA inter-strand crosslinks likely contribute to acetaldehyde mutagenesis

Amongst the adducts induced by acetaldehyde, inter- and intra-strand crosslinks have been shown to be highly mutagenic [10, 11, 14]. Pso2 in yeast has been demonstrated to be involved in repair of DNA ICLs induced upon psoralen and ultraviolet radiation exposure and *pso2* mutants demonstrate reduced mutation frequencies upon induction of DNA crosslinks [30]. We noted that the overall mutation frequencies in *Δpso2* yeast strains were lower than wild-type, indicating a role of Pso2 in bypassing endogenous DNA damage. We did not see any change in Can^R^ mutation frequencies in *Δpso2* yeast strains treated with 1% acetaldehyde for 1 hour as compared to the untreated controls. In addition, the increase in mutagenesis seen upon acetaldehyde treatment in the *Δrad1* strain was abolished in the *Δrad1Δpso2* double mutant strains (Fig 1C, Table S1). These data indicate that ICLs induced by acetaldehyde are repaired by nucleotide excision repair and require Pso2 activity for TLS and mutagenesis.

### Acetaldehyde-induced gCn→A mutations are elevated in *Δrad1* strains

We further obtained genomic DNA from Can^R^ yeast isolates either after treatment with acetaldehyde or after treatment with water (untreated) and sequenced their genomes. We analyzed 36 Can^R^ wild-type yeast strains treated with water, 36 Can^R^ wild-type yeast strains treated with acetaldehyde, 36 Can^R^ *Δrad1* isolates treated with water and 36 Can^R^ *Δrad1* isolates treated with acetaldehyde. Acetaldehyde-induced mutations were identified in the strains. Mutations common to two or more isolates were removed as those likely represented mutations preexisting in the cultures. Additionally, mutations in the parent strains were also removed. Analysis of the mutation spectrum in these isolates demonstrated that in both wild-type and *Δrad1* strains C→A changes were increased upon acetaldehyde treatment as compared to untreated strains (Fig 2A, B, Table S3). We then asked if the acetaldehyde-induced gCn→A mutation signature was elevated in the samples. We used the Trinucleotide Mutation Signature pipeline (TriMS) [19] to determine if gCn→A changes were enriched in the strains. TriMS compares the number of gCn→A changes to the number of C→A changes in the same data sets, as well as the number of cytosines and number of gcn motifs present in the background of each mutated residue. We saw a statistically significant enrichment of the gCn→A signature only in the *Δrad1* Can^R^ isolates obtained after treatment with acetaldehyde (Fig 2B, Table S3). The enrichment of the gCn→A signature strongly indicates that the increased mutation frequencies seen in the *Δrad1* isolates were due to an increase in acetaldehyde-specific mutagenesis.

**Figure 2.**
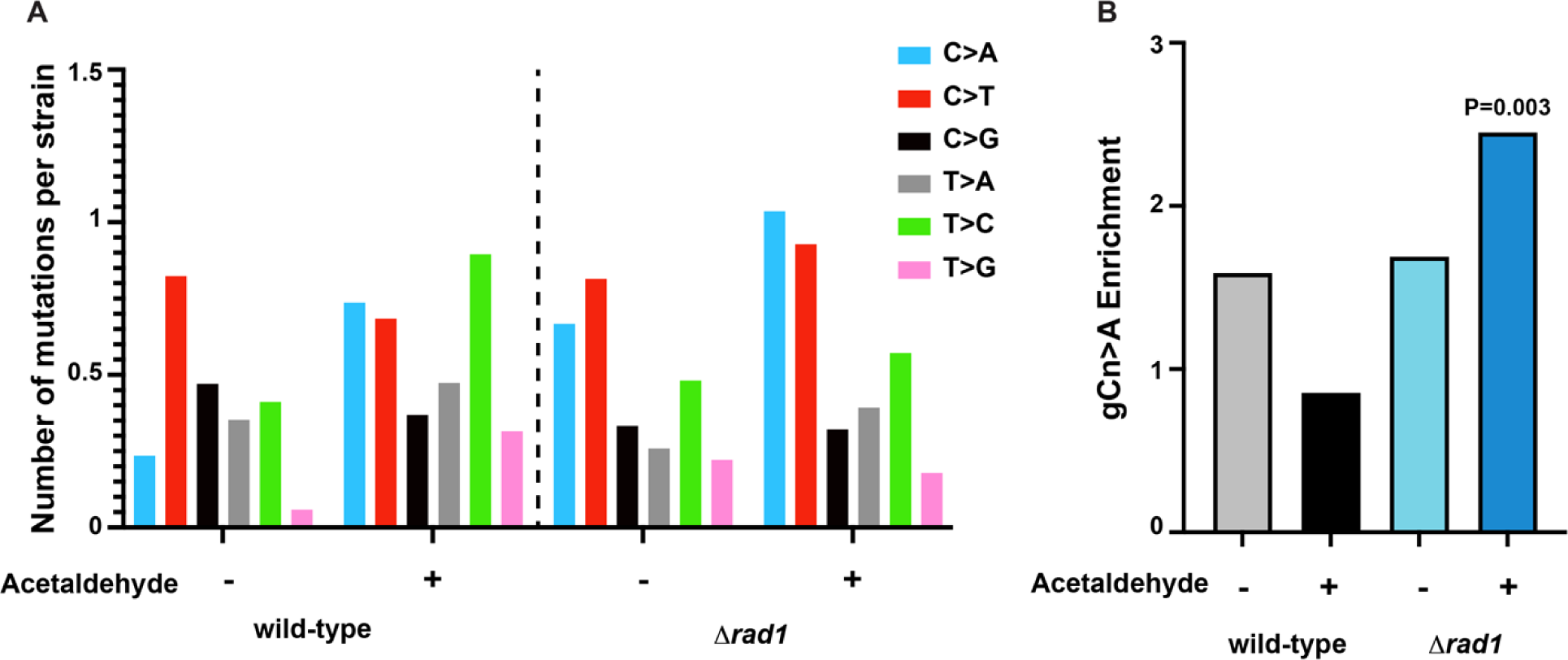
Acetaldehyde-induced mutation spectrum and signature in wild-type and *Δrad1* isolates. A) The number of single base substitution (given substitution along with substitution of the complementary base) per isolate with acetaldehyde or no mutagen control are shown. Mutation spectra is plotted as pyrimidine changes, taking into consideration reverse complements for each base change. B) Enrichment of the acetaldehyde-specific gCn→A mutation signature in the yeast strains. P-value was calculated in TriMS using a Fishers exact test as described in the methods.

### DPC repair defects increase acetaldehyde mutation frequencies

Wss1 (SPRTN in humans) and Ddi1 proteins are metalloproteases that have been shown to function in removal of formaldehyde-induced DNA-protein crosslinks (DPCs) [31-33]. To determine if the DPC repair pathway impacts acetaldehyde-induced mutagenesis in our strains, we deleted the genes *WSS1* and/or *DDI1* in yeast. We found that acetaldehyde-induced mutations frequencies were increased 1.5-fold in the *Δwss1* (P-value by a one-sided Mann Whitney U test = 0.0119) and 1.5-fold in *Δddi1* (P-value by a one-sided Mann Whitney U-test = 0.0043) isolates, each (Fig 3A). In the *Δwss1Δddi1* double mutant, we saw a further increase in acetaldehyde-induced mutation frequency (2.8-fold, P-value by a one-sided Mann Whitney U test = 0.0077) (Fig 3A, Table S1). We further noted that in contrast to the single deletion strains, the *Δwss1Δddi1* double mutant was highly sensitive to acetaldehyde, with a considerable loss of viability of the treated cultures (30%) compared to the untreated cells (Fig 3B, Table S2). Overall, we demonstrate that DPC repair is the second major pathway for prevention of acetaldehyde-induced mutagenesis and cell death.

**Figure 3.**
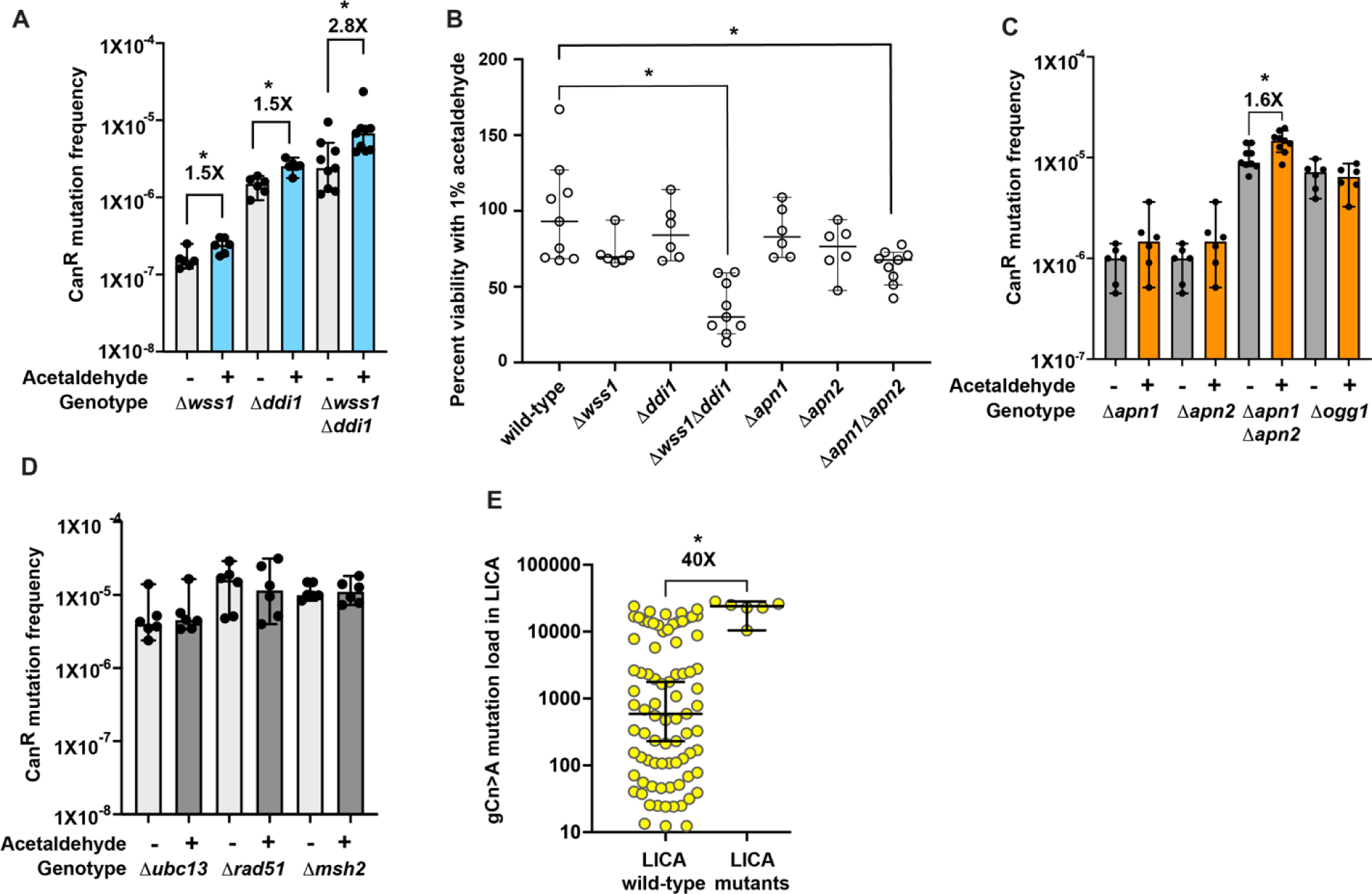
Multiple DNA repair pathways function to prevent acetaldehyde-induced mutagenesis. A) Mutation frequencies of yeast strains with defects in *WSS1* and/or *DDI1* mediated DPC repair. + indicates strains treated with 1% acetaldehyde and – indicates strains treated with no mutagen. Median mutation frequencies are plotted, error bars denote 95% confidence intervals. P-values were calculated using a one-sided Mann Whitney U-test. B) Quantitative survival assay of yeast strains with defects in DPC repair or deletion in *APN1* and/or *APN2* treated with acetaldehyde. Median and 95% confidence intervals are depicted. C) Mutation frequencies of BER defective yeast strains after treatment with acetaldehyde (+) or no mutagen (-). Median mutation frequencies are plotted, error bars denote 95% confidence intervals. D) Mutation frequencies of yeast strains defective in post-replicative repair (*Δubc13)*, homologous recombination (*Δrad51*) or mismatch repair (*Δmsh2*). Median mutation frequencies are plotted, error bars denote 95% confidence intervals. E) Analysis of gCn→A mutations in liver carcinoma samples from ICGC with either wild-type *ERCC4*, *SPRTN*, *FANCD2* (“wild-type”) or in samples with deleterious somatic mutations in the same genes (“mutants”). Only samples with statistically significant enrichment of gCn→A signature were analyzed. Mutation loads were calculated in TriMS as described in the methods. Median mutation frequencies are plotted, error bars denote 95% confidence intervals. P-values were calculated using a one-sided Mann Whitney U-test. * denotes P-values < 0.05.

### Other DNA repair pathways involved in preventing acetaldehyde-induced mutagenesis

Acetaldehyde exposure has also been linked to increased reactive oxygen species production (reviewed in [34, 35]). Such oxidative damage can lead to the accumulation of 8-oxo-guanine moieties leading to G→T/C→A mutations. 8-oxo-guanine residues are removed by the Ogg1 glycosylase and the resulting abasic sites are repaired via the activity of the BER enzymes Apn1 and Apn2 [36-39]. Deletion of *OGG1* and treatment with 1% acetaldehyde did not result in any increase in mutagenesis in yeast (Fig 3C, Table S1). On the other hand, while deletion of either *APN1* or *APN2*, individually, did not increase Can^R^ mutagenesis, we noted that the double *Δapn1Δapn2* mutants had slightly elevated mutation frequencies upon treatment with acetaldehyde (1.6-fold increase, P-value using a one-sided Mann Whitney U test = 0.0065) (Fig 3C, Table S1). In addition, we saw a decrease in cellular viability of the *Δapn1Δapn2* mutants to 67% upon treatment with acetaldehyde as compared to the wild-type isolates at 93% (P-value using Mann Whitney U test = 0.014) (Fig 3B, Table S2). As such, we conclude that while acetaldehyde does not appear to contribute to oxidative damage and accumulation of 8-oxo-guanine residues in our system, BER is likely involved in the repair of a minor subset of acetaldehyde lesions.

To test other DNA repair pathways that may be involved in repair of acetaldehyde-induced DNA damage in yeast, we deleted genes involved in post replication repair (*UBC13*) [40], mismatch repair (*MSH2*) [41] and homologous recombination (*RAD51*) [42], respectively. We did not see any increase in acetaldehyde-induced mutation frequencies in these strains (Fig 3D, Table S1).

### gCn→A mutations are higher in liver cancer samples with deleterious mutations in NER, DPC repair and FANCD2 genes

Previously, we demonstrated that liver, lung, head and neck and stomach cancers carry increased acetaldehyde-associated gCn→A mutations in the International Cancer Genome Consortium (ICGC) and the PanCancer Analysis WorkGroup datasets [19]. In these cohorts, we found that LICA (Liver Cancer) samples in ICGC had the largest number of tumors with statistically significant gCn→A enrichment (85/400). We further analyzed these samples to identify those tumors with somatic mutations in *ERCC4*, *ERCC1* (Rad1 and Rad10 endonucleases in yeast), *SPRTN* (Wss1 in yeast) and *FANCD2*. Rad10 works along with Rad1 in nucleotide excision repair [29], as such, both *ERCC4* and *ERCC1* were included in the analysis. Samples with *FANCD2* mutations were included because this gene has been previously shown to lead to increased mutations in *ALHD2^-/-^FANCD2^-/-^* mice, [7, 22, 25], indicating that this gene is involved in acetaldehyde-induced ICL repair. ICGC assigns the value of “high” impact to frameshift variants, non-conservative missense variants, initiator codon variants, stop gained and stop lost variants. Therefore, we used this criterion and identified four tumors with mutations in *ERCC4*, one tumor with mutation in *ERCC1*, two tumors with mutations in *FANCD2* of which one also had a mutation in *ERCC4* and one sample with a mutation in *SPRTN*. We further subset the number of tumors in LICA cohort to only analyze those samples that showed a positive and statistically significant enrichment of the gCn→A mutations. One sample with an *ERCC1* mutation was therefore removed since it did not have an enrichment of gCn→A mutations. We found that compared to tumors with wildtype *ERCC4*, *SPRTN* and *FANCD2*, the samples with mutations in any of these genes carried a higher gCn→A mutation load (wildtype samples, median = 591 gCn→A mutations per exome, mutant samples, median = 23,991 gCn→A mutations per exome) (Fig 3E, Table S4). Our data demonstrates that NER, ICL repair and DPC repair also function to prevent acetaldehyde-induced mutations in cancers.

## Discussion

In this study, we demonstrate that acetaldehyde-induced DNA damage is repaired by multiple pathways to prevent mutagenesis. Defects in NER, DPC repair, BER and ICL repair were found to alter acetaldehyde-induced mutation frequencies in yeast (Fig 4). Interestingly, similar to previous studies that showed that acetaldehyde was not mutagenic in DNA repair proficient human iPSCs, *Salmonella* strains, and yeast strains [17, 18, 28], we did not observe elevated acetaldehyde-induced mutagenesis in wild-type yeast strains, indicating that DNA repair pathways are highly efficient at removing acetaldehyde lesions in the genome.

**Figure 4.**
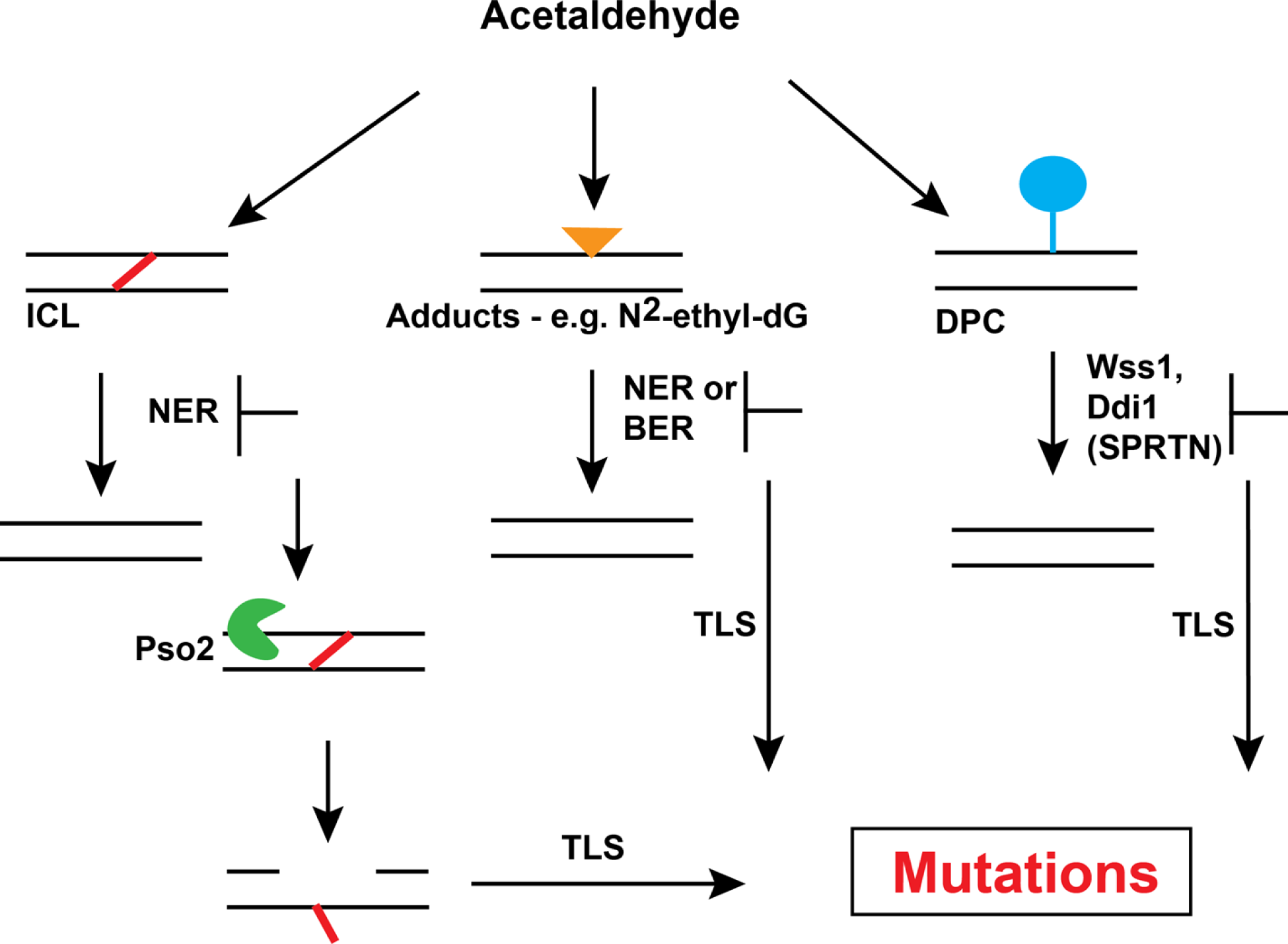
Model of DNA repair pathways that prevent acetaldehyde-induced mutagenesis. Acetaldehyde-induced inter-strand crosslinks (ICLs) shown in red, DNA adducts shown as a yellow triangle and DPCs shown as a blue circle connected to DNA. The different DNA repair enzymes and pathways that likely function to prevent or promote mutagenesis by acting on these adducts are shown.

The deployment of multiple DNA repair pathways to prevent acetaldehyde-induced mutagenesis signifies the induction of different types of DNA lesions generated upon acetaldehyde exposure in cells (Fig 4). Acetaldehyde is known to form 1,N^2^-propano-2′-deoxyguanosine [10-14, 34], and these lesions have been shown to block DNA replication [27, 43]. These guanine lesions are likely repaired via NER as such, deletion of *RAD1* or *RAD14* prevents excision of these lesions causing replication defects, recruitment of TLS and mutagenic bypass of the lesion.

NER has also been implicated in repair of ICLs (reviewed in [44]). Acetaldehyde also induces G-G ICLs. The motif gCn/nGc has two guanine residues on opposite strands near each other, wherein ICLs may be formed. As such, in the absence of *RAD1*, such ICLs likely persist in DNA and cause a replication block. *S. cerevisiae* predominantly uses Pso2 along with the helicase Hrq1 to endonucleolytically cleave DNA and unhook the ICL allowing for recruitment of a TLS polymerase and gap filling across the lesion [45]. In concordance with this model, we demonstrated that in *Δrad1Δpso2* strains acetaldehyde-induced mutagenesis is abrogated, likely due to the lack of TLS recruitment to the lesion.

Acetaldehyde exposure can also lead to the formation of DPCs which can further block DNA replication and lead to double strand breaks and genome instability. Studies using formaldehyde have demonstrated that DPCs are primarily repaired using the metalloproteases Wss1 (functional homolog of SPRTN) and Ddi1 in yeast. Deletion of both enzymes leads to extreme sensitivity of yeast to formaldehyde [31]. However, previously studies on DT40 cells showed that DPCs do not contribute to cytotoxicity by acetaldehyde [46]. On the other hand, work on *S. pombe* strains demonstrated that the Wss1 metalloprotease was responsible for DPC resolution and deletion of *WSS1* sensitizes cells to acetaldehyde exposure [27]. Here, our work demonstrates that both Wss1 and Ddi1 act redundantly to repair DPCs and prevent acetaldehyde-induced mutagenesis in yeast. Our finding was also corroborated in liver cancer cells where the sample with a deleterious mutation in SPRTN carried a much higher gCn→A mutation load than samples with wild-type SPRTN. Overall, our work and the studies in *S. pombe* together demonstrate that acetaldehyde-induced DPCs are a major source of genome instability.

Finally, previous reports have implicated ethanol consumption and acetaldehyde in generating oxidative stress. As such, we would anticipate that such acetaldehyde treatment would lead to an increase in 8-oxo-guanine levels which is predominantly repaired via BER. Unrepaired 8-oxo-guanines in DNA are bypassed via TLS and culminate in G→T/C→A mutations. We found that deletion of *APN1* and *APN2* in yeast led to a mild increase in acetaldehyde-induced mutagenesis. However, deletion of *OGG1*, the major glycosylase responsible for removal of 8-oxo-guanine, did not cause elevated acetaldehyde mutations. As such, we surmise that the gCn→A mutations are not a product of erroneous bypass of un-repaired 8-oxo-guanines. BER was also implicated in tolerance of acetaldehyde in *S. pombe [27]*. Based on these results, we propose that BER likely functions to remove DNA base adducts formed by acetaldehyde.

We further showed that whole exome sequenced liver cancer samples with deleterious somatic mutations in *ERCC4* (NER), *SPRTN* (DPC repair) and *FANCD2* (ICL repair) carry a higher burden of acetaldehyde-induced mutation signature as compared to tumors with no defects in these proteins indicating that the pathways modulating acetaldehyde-induced mutagenesis in yeast are conserved in humans.

Homologous recombination was shown to impact sensitivity of *S. pombe* to acetaldehyde [27] and CHO cells deficient in *RAD51D* were sensitive to acetaldehyde and demonstrated increased chromosomal aberrations [47]. However, we did not see increased acetaldehyde-induced mutation frequencies in *Δrad51* isolates. It is possible that in the absence of the NER, DPC repair and ICL repair pathways, homologous recombination or post replication repair pathways are required to prevent acetaldehyde-induced mutagenesis. Further, our assay is built to detect point mutations inactivating *CAN1*, as such, we do not know if these pathways are involved in preventing acetaldehyde-induced double strand breaks and gross-chromosomal rearrangements. Further investigations are justified to determine the role of homologous recombination pathways in maintaining genome stability upon acetaldehyde exposure.

Acetaldehyde is a known carcinogen and a DNA damaging agent. We previously showed that an acetaldehyde-specific mutation signature can be detected in cancers, indicating that it is a source of DNA damage and mutagenesis in tumors [19]. As such, understanding the mechanisms that contribute to it is essential to determine if certain individuals may be predisposed to increased genome instability upon acetaldehyde exposure. We demonstrated here that defects in a variety of DNA repair pathways can lead to elevated mutagenesis upon acetaldehyde exposure in yeast and in liver cancers demonstrating the various types of DNA damage induced by acetaldehyde exposure. Considering that acetaldehyde is both an endogenous and environmental carcinogen, understanding the molecular mechanisms that alter acetaldehyde mutation rates is essential to enable determination of “at-risk” individuals susceptible to acetaldehyde-induced cancers.

## Methods

### Strains

Yeast strains used in this study are derived from the ySR128 (*MATα ura3Δ can1Δ ade2Δ leu2- 3,112 trp1-289 lys2:ADE2-URA3-CAN1).* Deletions of the genes *RAD1, RAD14, APN2, WSS1, DDI1, PSO2, REV3 and OGG1* using either *KANMX* or *HPHMX* cassettes, conferring resistance to G418 and hygromycin, respectively. *APN1* was deleted using the *bsd* gene conferring resistance to blasticidin. The complete strain list is provided in Table S5.

### Acetaldehyde treatment and mutation frequency measurement

Yeast strains were incubated at 30°C shaking at 160RPM overnight. The next day, the yeast cells were counted on a hemocytometer and diluted 1:10 in 30ml of fresh media to yield a starting concentration of 1×10^7^ cells/ml. The 30ml yeast cultures were incubated at 30°C with shaking at 160 RPM for three hours. The cultures were then spun in 50ml conical tubes at 2500 RPM for five minutes. The cultures were washed once with water and then split into two 15ml samples in 15ml conical tubes. One set of the 15ml conical tubes contained yeast cells treated with water. The second set of 15ml conical tubes contained yeast cells with 1% acetaldehyde (Sigma, catalog number 402788) diluted in water. Acetaldehyde was kept chilled and chilled pipet tips were used to prevent evaporation. Both sets of tubes were incubated for one hour at 30°C in a rotary shaker. The cultures were then spun down and washed once with water. The cultures were resuspended in 2ml of water. Appropriate dilutions of cells were plated on complete synthetic complete (SC) media (MP Biomedicals) to measure viability and SC-Arginine plates containing 60mg/ml canavanine (Sigma). Plates were incubated at 30°C for five days.

Colonies were counted using the aCOLyte 3 automated colony counter (Synbiosis). The mutation frequency was calculated using the formula –

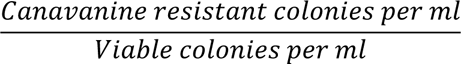

Cellular viability after acetaldehyde treatment was calculated as –

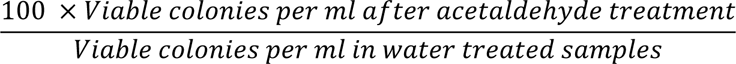

### Whole genome and mutation analysis

Individual Can^R^ isolates were obtained for wild-type and *Δrad1* isolates after treatment with either water or with 1% acetaldehyde. The colonies were streaked out to obtain pure cultures and genomic DNA was extracted using the Zymo YeastStar genomic DNA isolation kit (Genesee Scientific). 10ng/ul DNA was used for library preparation with the Watchmaker DNA library prep with fragmentation (Watchmaker Genomics). Each sample was provided a unique dual index adapter (stubby adapter IDT, xGen UDI 10nt Primer Plate 1-4 IDT). Samples were pooled and sequenced on the 1 lane of Illumina NovaSeq 6000 S4 PE 2×150 sequencing system. The resulting FASTQ files were aligned to the reference genome ySR128 [48] using BWA mem [49], duplicate reads were removed using Picard tools (http://broadinstitute.github.io/picard/) and mutations were called using VarScan2 using a variant allele frequency filter of 90% [50-52]. The untreated original cultures were also sequenced. Mutations present in the untreated cultures as well as mutations common to two or more samples were marked as pre-existing and removed.

TRInucleotide Mutation Signatures (TriMS) [19] was used to further measure enrichment of the gCn→A mutation signature in the acetaldehyde and water-treated isolates. Specifically, TriMS compares the total number of gCn→A mutations in a sample to the number of C→A changes in the same sample, as well as the number of cytosines and gcn motifs within 20bp of a mutated residue. An enrichment greater than 1 is further tested statistically using a one-sided Fishers’ exact test. The code for TRIMS is deposited in github and can be accessed at https://github.com/nataliesaini11/TriMS.

### Mutation analysis in cancers

Whole exome sequenced tumors in LICA-CN cohort from ICGC were analyzed using TRIMS as described in [19]. We further determined the samples carrying deleterious defects in either *ERCC1, ERCC4, FANCD2* and *SPRTN* via the ICGC data portal - https://dcc.icgc.org/. The mutation loads attributable to acetaldehyde-induced gCnàA changes was calculated by TRIMS using the equation –

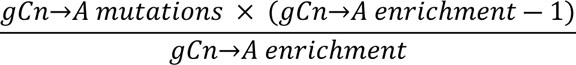

## Supplementary Tables

**Table S1 –** Data table for Figures 1B, 1C, 3A, 3C and 3D showing *Can^R^* mutation frequencies in yeast treated with acetaldehyde.

**Table S2 –** Source data for figures 1D and 3B showing viability of strains treated with acetaldehyde as compared to untreated strains.

**Table S3 –** Data for Figure 2. Whole genome sequencing data from yeast strains in this study. S3A - Mutation list from whole genome sequenced yeast strains; S3B - Mutation spectrum of acetaldehyde and water-treated isolates; S3C – gCn→A mutation signature analysis using TriMS.

**Table S4 –** Source data for Fig 3E showing gCnàA enrichment and mutation loads in liver cancer samples (LICA) with and without somatic mutations in *ERCC4*, *SPRTN*, *FANCD2*.

**Table S5 –** Yeast strains used in this study.

## Data Availability

The yeast strains used in the study are available upon request. Raw FASTQ sequence files from whole-genome sequencing of yeast samples are being deposited to the Sequence Read Archives (SRA) database. Sequence for the reference yeast genome used in this study (ySR128) is accessible on SRA under PRJNA524644. Source code for TriMS is available on GitHub (https://github.com/nataliesaini11/TriMS).

## Acknowledgements

We would like to thank Dr. Judit Jiminez Sainz and Mr. Thomas Blouin for reading the manuscript and providing helpful comments.

